# Sensory prediction errors, not performance errors, update memories in visuomotor adaptation

**DOI:** 10.1101/313551

**Authors:** Kangwoo Lee, Youngmin Oh, Jun Izawa, Nicolas Schweighofer

**Affiliations:** Biomedical Engineering, University of Southern California, Los Angeles, USA; Neuroscience, University of Southern California, Los Angeles, USA; Empowerment Informatics, University of Tsukuba, Tokyo, Japan; Biokinesiology and Physical Therapy, University of Southern California, Los Angeles, USA

## Abstract

Sensory prediction errors are thought to update memories in motor adaptation, but the role of performance errors is largely unknown. To dissociate these errors, we manipulated visual feedback during fast shooting movements under visuomotor rotation. Participants were instructed to strategically correct for performance errors by shooting to a neighboring target in one of four conditions: following the movement onset, the main target, the neighboring target, both targets, or none of the targets disappeared. Participants in all conditions experienced a drift away from the main target following the strategy. In conditions where the main target was shown, participants often tried to minimize performance errors caused by the drift by generating corrective movements. However, despite differences in performance during adaptation between conditions, memory decay in a delayed washout block was indistinguishable between conditions. Our results thus suggest that, in visuomotor adaptation, sensory predictions errors, but not performance errors, update the slow, temporally stable, component of motor memory.

## Introduction

What is the mechanism that enables adaptation in response to a sensory perturbation? A long-held theoretical view is that sensory-motor adaptation is due to the update of internal forward models ^1,2^, which are neural processes that predict the sensory consequences of motor commands ^3–5^. According to this view, when a sensory perturbation such as a visuomotor rotation alters the sensory outcome, the forward model is updated to minimize the sensory prediction error (SPE), that is, the error between the sensory outcome and its prediction ^6^. Forward model update can happen continuously during daily activities via the individual’s own movements (“self-supervised learning”;^3,7^). The output from the updated forward model can then be used to compute the motor command needed to compensate for the perturbation when a task goal is defined ^8^.

Thus, in theory, SPEs alone in the absence of a task goal are sufficient for motor adaptation via forward model update. Decisively, movements during adaptation do not need to be goal-directed because only the difference between actual and predicted sensory consequences matters in the update of the forward model ^9,10^. However, in most real-life activities and in most behavioural experiments, performance errors or rewards, whether explicit (e.g., score, evaluation) or implicit (e.g., self-evaluation), related to the goal are available. Because reward-related signals have been shown to influence motor adaptation ^8,11–14^ both SPEs and performance errors could simultaneously play a role in adaptation.

A seminal experiment by Mazzoni and Krakauer ^15^ suggested that SPEs in the absence of initial performance errors lead to a drift in hand direction. In this experiment, participants made movements to one of eight targets around a circle, spaced 45 degrees apart. A 45-degree visuomotor perturbation was then suddenly introduced, creating large performance errors between the target and the cursor. After the second adaptation trial, participants were told that shooting at the neighboring target placed clockwise of the main target would cancel the error. Indeed, in the trials following the introduction of this strategy, the performance error between the main target and the cursor was approximately zero. However, a drift appeared in the later trials, with the cursor starting to rotate clockwise from the main target. After about 70 trials, participants were told to stop using the strategy and aim directly at the main target again. The following trials showed strong after-effects. These results suggest that the SPE, but not the performance error between the cursor and the main target, is involved in adaptation via updates of an internal model. This is because just after the strategy is introduced, the drift appears despite zero performance error.

However, this experiment did not allow for a disambiguation of the roles of SPEs versus performance errors on updating motor memories, because both types of errors were present in the drift phase. An extension of this initial study by Taylor and Ivry ^16^ with a larger number of adaptation trials showed that the performance error between the cursor and the main target that appears during the drift phase influences performance during adaptation: when the number of trials increased, participants started to generate corrective movements to cancel the drift. In addition, a second performance error, the error between the cursor and the neighboring target, which corresponds to the aiming error, could also influence adaptation. Note that this second performance error would initially act on the drift in the same direction as the SPEs. We call the first type of performance error, the error between the main target and the cursor, PE1 (See Figure 1A). We call the second type of performance error, the error between the neighboring target and the cursor, PE2 (See Figure 1A).

**Figure 1.**
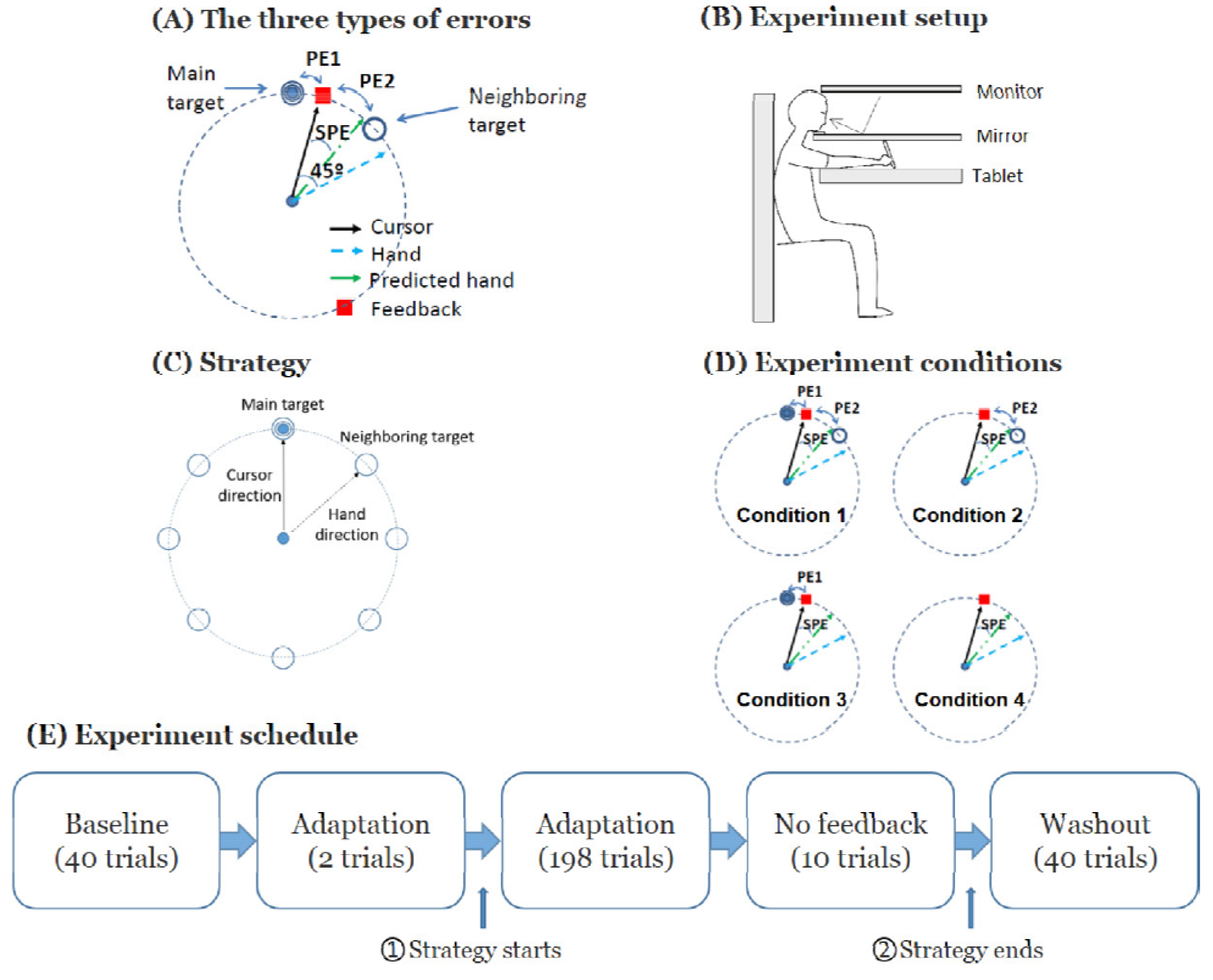
Experimental design to manipulate the influence of possible types of error on visual motor adaptation. (A) The three possible types of error influencing performance and adaptation. (B) The experimental set-up for visuomotor motor adaptation. We acknowledge Cheolhwan Sim for drawing this figure. (C) Strategy given to the participants to counteract the perturbation: participants were told to shoot to the neighboring target in order to reach the main target. (D) The four experimental conditions. In Condition 1, both the main and the neighboring target remained on the screen following the shooting movement. In Conditions 2 and 3, the main target or theneighboring target disappears as the hand was at the 1/3^rd^ distance between the home position and the target circles, respectively. In Condition 4, both targets disappeared. (E) Experiment schedule common to all conditions. The explicit strategy was given at time ⍰, and the participants were asked to stop using the strategy from time ⍰.

Here, we extended the study by Taylor and Ivry ^16^ in two ways: First, we performed a two-by-two design experiment to distinguish the possible influences of SPE, PE1, and PE2 on adaptation: the main target, the neighboring target, both targets, or none disappear following the movement onset (Figure 1A). When no targets disappear (Condition 1), the learner can rely on PE1, PE2, and SPE. When the main target disappears (Condition 2), the learner can only reliably rely on PE2 and SPE. When the neighboring target disappears (Condition 3), the learner can only reliably rely on PE1 and SPE. When the both the main and the neighboring target disappear (Condition 4), estimation of both PE1 and PE2 becomes unreliable, and if adaptation occurs, we hypothesized that it is driven by SPE. Second, we tested after-effects in a delayed washout block, given after a no-feedback movement block in all four conditions, to probe whether the slow component of memory ^17,18^ is updated via performance errors or SPEs.

Our results show that participants in the two conditions with PE1 performance errors clearly improved performance during adaptation by reducing the magnitude of the drift, and exhibited smaller drifts than participants in the two conditions without PE1. However, in delayed washout, both the amplitude and the decay were not distinguishable in the four different conditions, despite differences in performance during adaptation. Thus, our results support the view that the slow, temporally stable, component of visuomotor adaptation in arm movements is updated by SPEs, not performance errors.

## Materials and methods

### Detailed Experimental methods

#### Participants

Fifty-two young and right-handed participants (23.4±0.6 years old, 35 females; all results are reported as mean ± standard errors) with no history of neurological disorders participated in this study. We randomly assigned the participants to one of four conditions (N=13 per condition). All participants signed an informed consent form and were right-handed based on the Edinburgh handedness inventory. The study was approved by the Institutional Review Board at the University of Southern California (HS-08-00461) and was performed in accordance with the approved guidelines.

#### Set-up

Participants were asked to sit on a fixed chair in front of a horizontal double-layered device (Figure 1B). The device has a layer with a reflected monitor (visual space) and a tablet layer (hand space). Participants were instructed to move a digitizer pen (Wacom Intuos 7) on the tablet. When the lights were turned off in the experimental room, the participants’ view of their forearm and hand was obscured by the mirror. Head and trunk movements were minimized by using a chin-rest and a fixed chair. A cursor (0.1 cm radius) representing the tip of the pen was displayed on the mirror.

#### Timing of a single trial

Before the start of each trial, a red circle was displayed on the reflected screen with a radius equal to the distance between the cursor and a white home circle (2.6 mm radius). Participants were instructed to move the digitizer pen to the white home circle by minimizing the radius of the red circle. At the onset of the trial, a cursor (red dot with 1 mm radius, whose position matched the tip of the digitizer pen) was displayed, and a main target (bull’s eye shape) appeared at one of eight possible locations on a circle of 5.3 cm radius. Target locations were distributed evenly on the circle with 45 degrees between targets (except in the no-feedback block where no target and no feedback were shown). At each trial, the location of the main target was pseudo-random, with each target being presented once for each block of eight targets. Upon presentation of the main target, shown as a bull’s eye, participants were asked to perform an outward shooting movement passing the target circle. The cursor disappeared when the distance between the home circle and the cursor was greater than one-third of the circle radius. A 4.2mm sized red square was displayed on the target circle to indicate where the cursor has passed. Different types of visual feedback were given in different blocks and conditions (as discussed below). After each shooting movement, participants were instructed to perform an inward movement to the white home circle as before the start of each trial. When the duration of the shooting movement was outside the 50 to 300ms range, the messages ‘Too fast’ or ‘Too slow’ was displayed.

#### Overall design for common to all conditions

The experiment schedule consisted of a baseline block, an adaptation block, a no feedback block, and a washout block (Figure 1F). Before the experiment, participants participated in a familiarization block without visuomotor rotation of 110 trials. Following familiarization, they performed a baseline block of 40 trials with no perturbation. Following baseline, a 45 degrees counter-clockwise adaptation was introduced (a small amount of noise, 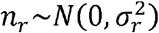, σ_*r*_ = 0.05 degree was introduced; pilot data, showed this noise slowed down adaptation slightly, possibly because it reduces the effectiveness of cognitive strategies. Note, however, that this noise was small compared to the trial-by-trial standard deviation of 6.1 degrees in average for all participants at baseline). During the adaptation block, two targets were presented at each trial: a main (bull’s eye) target and a neighboring target, shown at 45 degrees clockwise of the main target. As in Mazzoni and Krakauer’s work ^15^, participants were told about the rotation and were then instructed to use the following strategy after the second adaptation: “Counter the rotation by aiming at the clockwise neighboring target rather than at bull’s eye” (Figure 1C).

After 198 trials of adaptation trials with this strategy, participants performed a block of 10 shooting trials without feedback before performing a delayed washout block. The main goal of this no-feedback block was to erase the fast component of adaptation ^17,18^, and therefore to test in the delayed washout block whether the slow component of adaptation had been updated via performance errors, SPEs, or both. During this no-feedback block, participants were instructed to shoot toward an arbitrary direction on a 180-degree arc displayed ahead of the home position, with the same radius as the target circle in the other blocks, and to return to the home position. No feedback was presented during and after the shooting movements.

We expected that the fast component of adaptation would be fully erased following these blocks for two reasons. First, whereas the block lasted 154 s on average, the time constant of the decay of the fast component of visuomotor adaptation has been estimated to be approximately 16.5s ^19^. Second, because participants performed active movements, forgetting of the fast component is faster than during an equivalent rest period ^20,21^. Following this no-feedback block, participants were then instructed to stop using the explicit strategy and to shoot directly at the main target. The 80 trials following these new instructions formed the delayed washout block.

#### Experimental conditions

During the adaptation block, extending the method of Taylor and Ivry ^16^, we manipulated the possible contributions of PE1 and PE2, by removing the bull’s eye target, the neighboring target, or both, as soon as the cursor passed 1/3rd of the distance between the center of the home position and the target circle (corresponding to 1.77 cm away from the center home position). When the cursor re-appeared to show where it crossed the circle, both PE1 and PE2 were shown (Condition 1), only PE1 was shown (Condition 2), only PE2 was shown (Condition 3), or neither was shown (Condition 4) - see Figure 1E.

#### Statistical analysis

The angular error was computed as the difference between the main target angle and the shooting angle. The shooting angle was given by the line connecting the center position to the point on the invisible target circle given by interpolation of the two closest data points. To assess differences in adaptation and delayed retention among the four conditions, we performed one-way ANOVAs on the mean adaptation angular error of the middle 10 adaptation trials, last 10 adaptation trials, and first 10 washout trials in each condition.

To estimate the amplitude and rate of adaptation and washout, we then modeled the adaptation data and washout data with single exponential models of the form, *Err(t)* = *A exp (− t/ τ)*, where *A* is an amplitude parameter and *τ* a time constant parameter. Estimates and confidence intervals for *A* and *τ* and for the goodness of fit R^2^ were obtained with a bootstrap procedure with 10000 samples by drawing participants with replacement.

Finally, to test whether the performance at the end of adaptation predicted performance at the beginning of washout, we correlated the average performance in the last 10 trials of adaptation and the first 10 trials of washout.

## Results

### Adaptation block

Figure 2 shows the performance error between the main target and the cursor for representative participants in each condition. Figure 3 shows the average error across participants. In all conditions, the strategy was effective at canceling the perturbation initially during the adaptation phase but participants subsequently exhibited a drift, as shown previously ^15,16^. However, when PE1 was present, participants often attempted to compensate for the drift by generating corrections to reduce the performance error between the main target and the cursor (see Figure 2). Such corrections, which were highly variable both within and between participants, resulted in smaller drifts overall (Figure 3; compare Conditions 1 and 3 to Conditions 2 and 4). ANOVA and post-hoc t-tests confirmed this reduction in drifts during adaptation in conditions with PE1 present (Figure 4(A); between-condition differences in error in mid and last 10 trials: ANOVAs *p* = 0.0001 and *p* = 0.003, respectively). The drift in the 10 middle adaptation trials of Condition 1 (with PE1 and PE2) was smaller than that in Condition 2 (PE2 only) and Condition 4 (neither PE1 nor PE2) (all *p* < 0.05; Tukey test). The drifts in the last 10 adaptation trials of Conditions 1 and 3 were smaller than in Conditions 2 and 4 (all *p* < 0.05; Tukey tests). However, the final level of adaptation was not different in Condition 4 (no feedback) from that of Condition 2 (with PE2, difference in error in middle and end of adaptation in Conditions 2 and 4, both *p* > 0.5). Thus, the presence of PE2 did not induce greater drift. Exponential fits to mean adaptation data (Figure 5A) shows overall group similarities between adaptation in Conditions 1 and 3, on one hand, and Conditions 2 and 4, on the other hand. We note that the data of Condition 4 shows that neither PE1 nor PE2 was needed for the drift to be induced. This result is consistent with the drift being induced by the SPE only.

**Figure 2.**
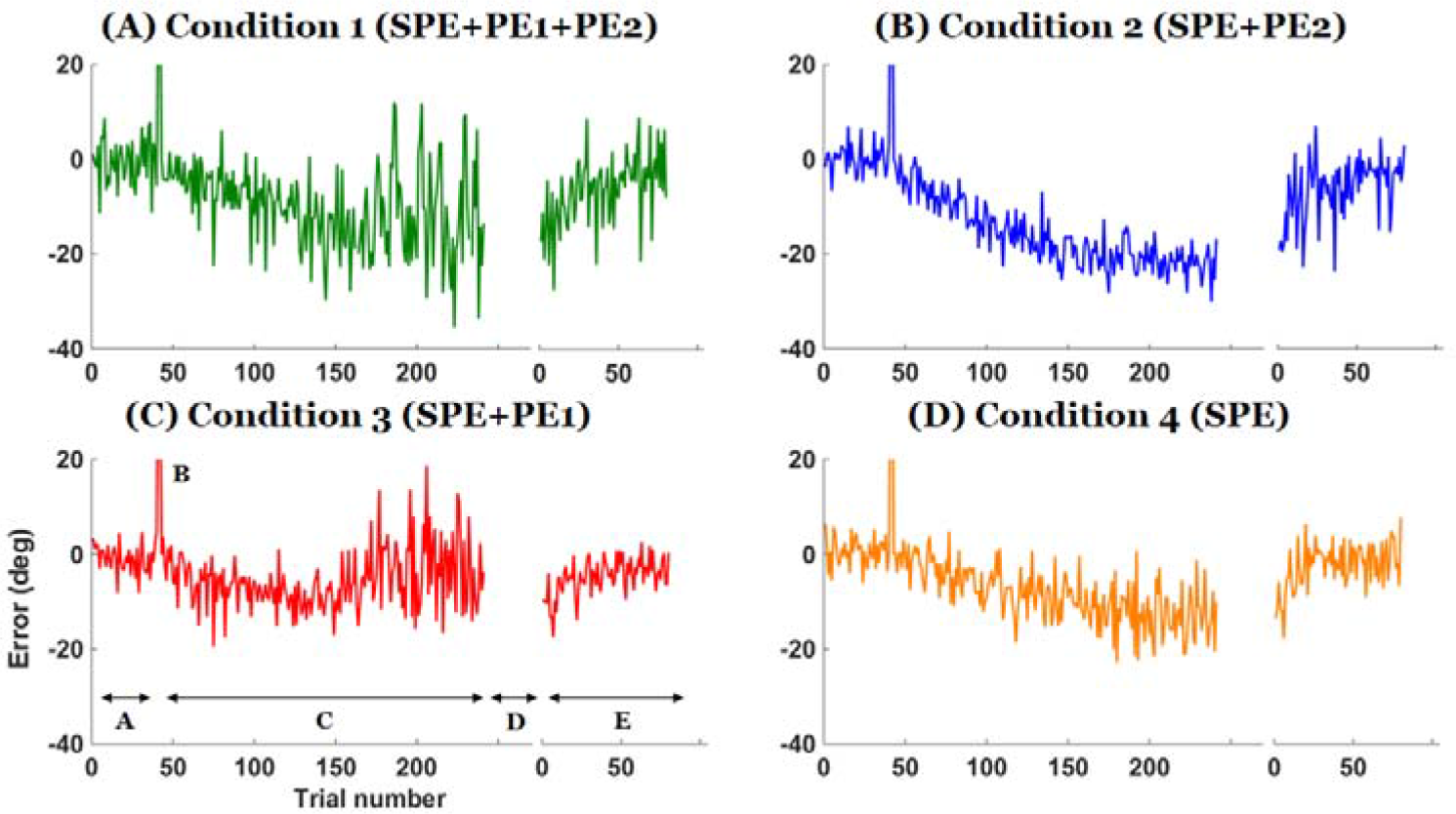
Representative individual data of the error between the main target and the cursor at the end of the movements in the four experimental conditions. Phases of the experiments are shown in bottom left panel. Phase A is the baseline block, B shows the two trials of perturbation without strategy, C is the adaptation block with strategy, D is the no-feedback block, and E is the washout block. Note how the strategy is effective in canceling the perturbation initially, how a drift appears in all conditions, and how attempts to minimize the error between the main target and the cursor in Conditions 1 and 3 reduce the drift overall; however, participants show strong after-effects when the strategy is removed in all conditions.

**Figure 3.**
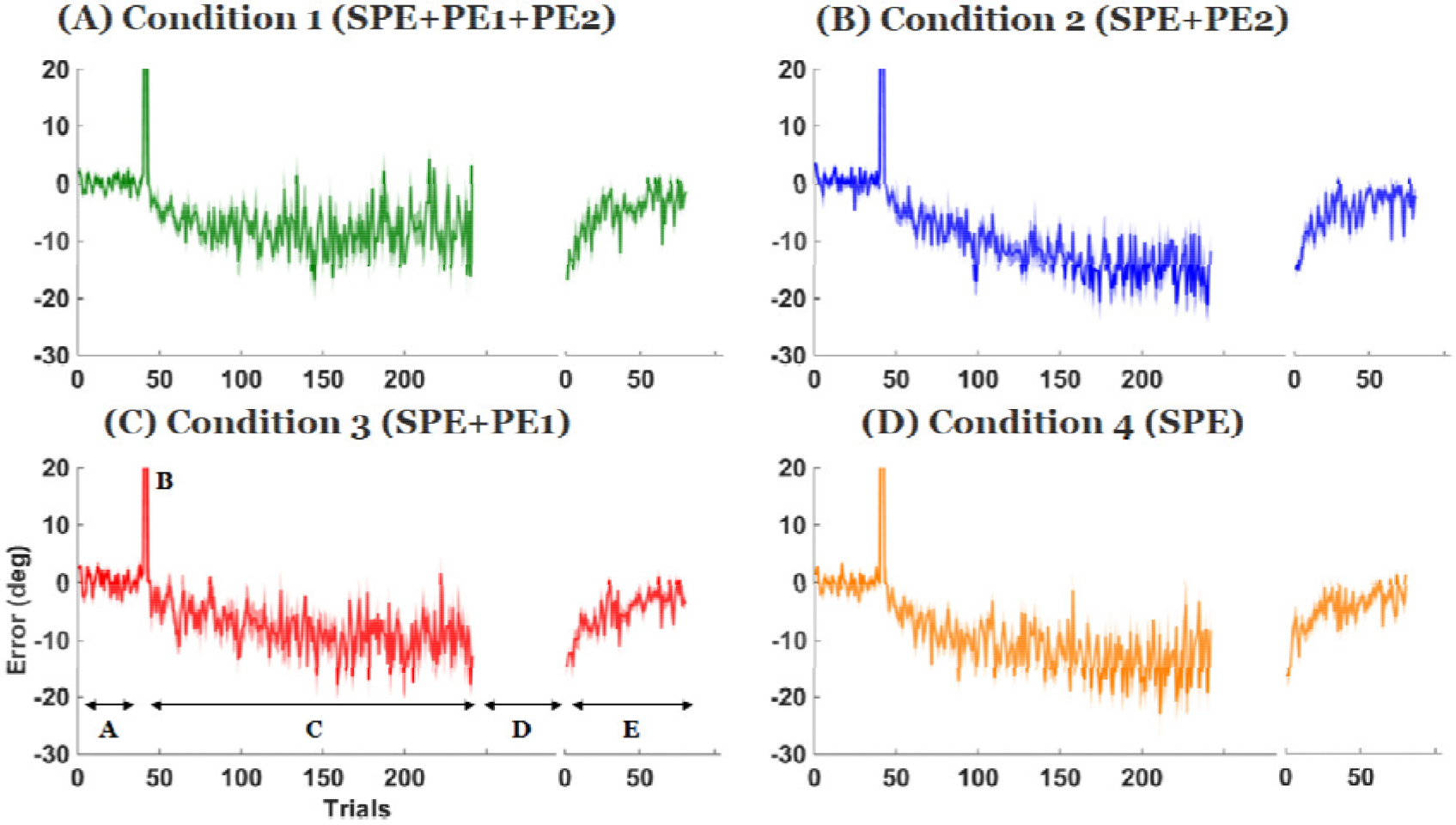
Mean error between the main target and the cursor at the end of the movements. Each phase represents the same blocks as in Figure 2. Shaded area represents the standard error across participants at each trial. Note how attempts to minimize the drift in Conditions 1 and 3 reduce the drift overall; however, participants exhibit similar washout in all conditions.

**Figure 4.**
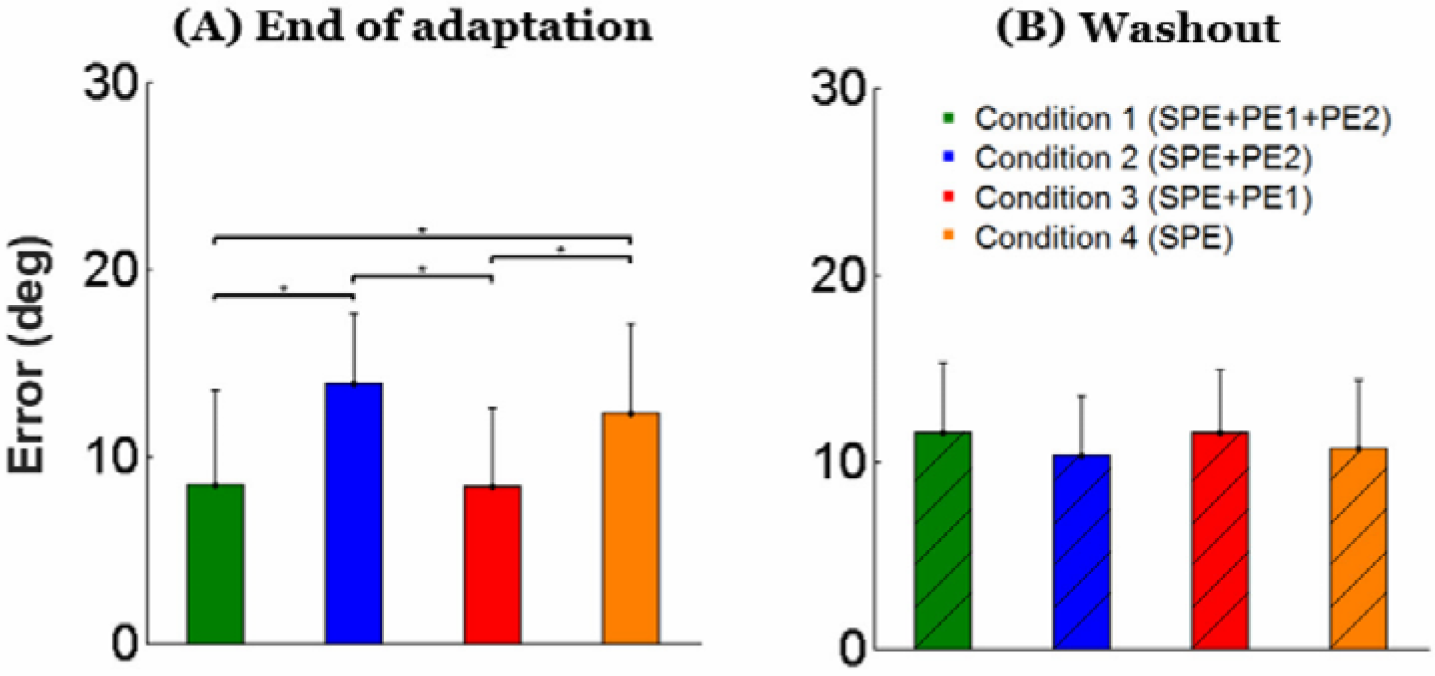
Mean angular error between main target and cursor in each block in the end of adaptation (A), and in washout (B) in all conditions.

**Figure 5.**
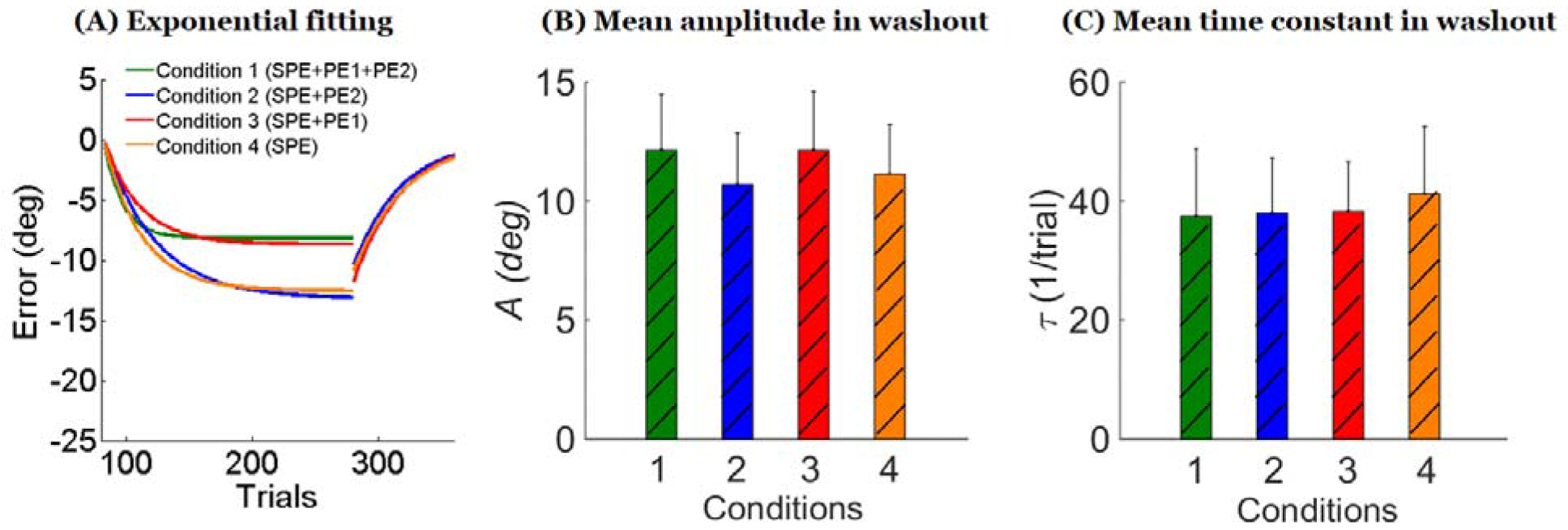
Exponential fits to adaptation and washout data. (A) Exponential fits for each group for both the adaptation phase and the washout phase. Note how the exponential fits during adaptation blocks are largely different between conditions with and without PE1, but the fits in washout are strikingly similar in all conditions. (B) Amplitude parameter *A* of the exponential decay in washout estimated via a bootstrap analysis shows no differences between conditions (mean and 95% confidence interval, see text). (C) Time constant τ of exponential decay in washout (mean and 95% confidence interval).

We note that the exponential fit during the adaptation phase is relatively good in Conditions 2 and 4 (R^2^ estimated from bootstrap = 0.44 ± 0.11SD, and 0.29 ± 0.12, respectively) was but poor in Conditions 1 and 3 (R^2^ = 0.10 ± 0.05, and 0.16 ± 0.10). This suggests that performance in Conditions 2 and 4, was overall decreasing with an exponential-like shape, with relatively low noise. In contrast, in Conditions 1 and 3, high noise and corrections towards baseline, as seen in Figures 2 and 3, created performance patterns that were not well fit by exponential functions.

### Delayed washout block

As seen in Figure 3, the after-effects in the washout periods were strikingly similar between conditions. Indeed, the mean of the first 10 trials of the washout was not different across conditions (ANOVA *p* > 0.5). Exponential curves fitted of the average washout data in each condition almost perfectly overlapped in all conditions (Figure 5A). This similarity between decay during washout in the four conditions was verified by a bootstrap analysis. In contrast to the adaptation phase were exponential fit was poor for Conditions 1 and 3, performance during the washout phase was relatively well fit by exponential functions in all conditions (R^2^ estimated from bootstrap: Condition 1: R^2^ = 0. 48 ± 0. 05 SD, Condition 2: R^2^ = 0. 44 ± 0.07; Condition 3: R^2^ = 0. 54± 0.07, and Condition 4: R^2^ = 0. 49 ± 0.07). Figure 5B and 5C shows that the 95% confidence intervals for the exponential amplitudes and time constants *τ* derived from the bootstrap analysis largely overlapped in all conditions.

Performance at the end of adaptation predicted after-effects during washout in conditions without PE1 (Conditions 2 and 4), but not in conditions with PE1 (Conditions 1 and 3). This is shown by significant correlations between the average performance in the last 10 trials of adaptation and the first 10 trials of washout in Conditions 2 and 4 (*p* = 0.02, and *p* = 0.005), but not in Conditions 1 and 3 (*p* = 0.14 and *p* = 0.32), as shown in Figure 6.

**Figure 6.**
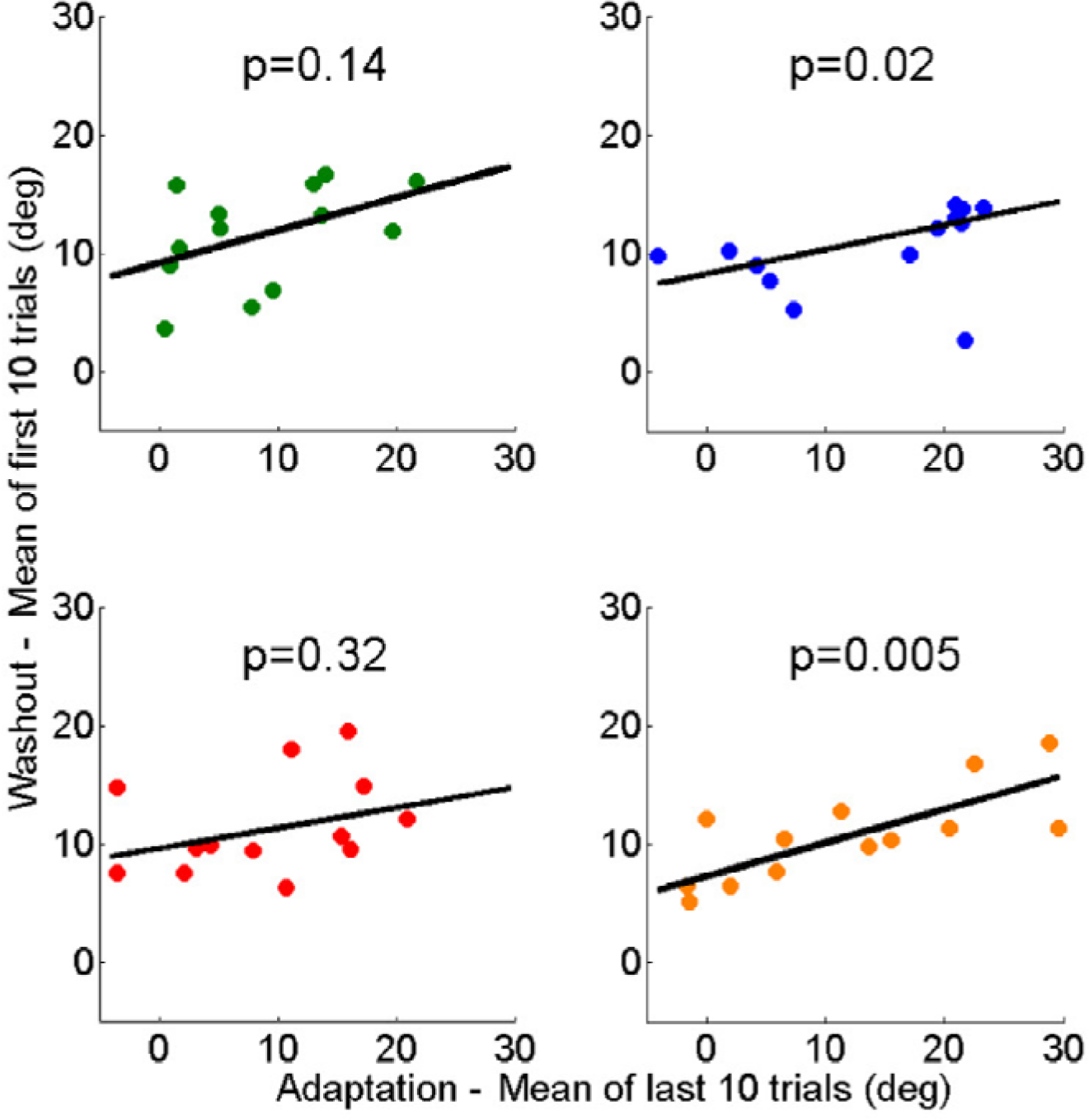
End of adaptation versus washout: regression of the first 10 trials of washout as a function of the last 10 trials of adaptation. Performance at the end of adaptation predicted performance in washout block in conditions without PE1 (Condition 2: *p* = 0.02 and Condition 4: *p* = 0.0005), but not in conditions with PE1 (Condition 1: *p* = 0.14 and Condition 3: *p* = 0.32).

In sum, conditions with PE1 (Conditions 1 and 3) showed a prominent difference between performance during adaptation and performance in retention, as probed by the delayed washout test. In contrast, in conditions without PE1 (Conditions 2 and 4), performance during adaptation predicted washout. In washout, there was no between-subjects difference in both mean performance and variability around this mean.

## Discussion

In this study, we manipulated the availability of performance errors following fast shooting movements in a visuomotor adaptation experiment in which participants were instructed to correct the given perturbation by strategically shooting to a neighboring target. Our experimental results show the followings. First, we replicated the drift following the introduction of the strategy, as found previously in Mazzoni and Krakauer ^15^, and Taylor and Ivry ^16^. The drift appears very robust and implicit, as it was shown in all conditions despite clear instructions to minimize the error between the cursor and the target. Second, the drift was reduced by the performance errors between the main target and the neighboring target when available following the movement (PE1): Results from Conditions 1 and 3 show that, as the number of trials increased, participants reduced the drift. Note that in Taylor and Ivry ^16^, in what is equivalent to our Condition 1 but with 322 adaptation trials, the average drift became near zero as the number of trials became large. In contrast, with 200 trials, we found that such reduction of drift is incomplete in amplitude (participants did not completely cancel the drift, even at the end of adaptation) and in consistency, as there was a large variability in drift reduction both within and between participants. Third, comparing performance results of Conditions 2 and 4 shows that the drift was not influenced by the second possible performance error, PE2, between the cursor and the neighboring target. This is an important control to the determination of the role of SPEs in adaptation with this paradigm, because PE2 could have initially generated the drift as it is in the same direction as the SPEs. Thus, only the performance error PE1 had an effect on behavior during the adaptation phase, by minimizing the drift via corrective movements towards the main target. Fourth, our results of Condition 4 show that participants exhibited the drift without target error feedback. This result is in line with previous studies showing visuomotor adaptation without explicit targets ^9,10^; but see Gaveau and Prablanc ^22^. Fifth, despite large differences in performance during adaptation between conditions with and without PE1 available, there was no difference between conditions in delayed washout, with a remarkable superimposition of the decay in washout across conditions.

The most parsimonious explanation of these results is that SPEs update the motor memory, and that such updates do not depend on the actual motor commands during adaptation. The drift can be best explained by update of the memory from SPEs alone because initially, the amplitude of SPE was near 45 degrees, but the amplitude of the performance error PE1 was near zero. As the drift increased, so did the amplitude of PE1, however. Because after-effects in the delayed washout blocks were highly consistent despite large difference in performance during adaptation (see Figure 4A), the actual performance during adaptation had no, or little, role in updating the temporally stable motor memories; this shows that the update of the internal estimate of the perturbation does not depend on the motor commands. In contrast, performance during adaptation was modulated by the strategy, the drift, and especially by the presence of PE1, via corrections aimed at counteracting the drift. How can the update of the temporally stable component of adaptation be independent of performance during training since the forward models receive, by definition, the motor command as input? The SPE is the difference from actual sensory feedback and predicted feedback, which both depend on the motor command; thus, in this subtraction, the effect of the motor command on actual sensory feedback is canceled by the effect of the motor command on predicted sensory feedback. As a result, internal model update happens identically independently of the actual movements in the adaptation phase.

To better understand this argument, we present here a simple model of visuomotor-adaptation: On trial *t*, the learner generates a motor command *u*_*t*_ to reach a target *t*_*t*_. Here, we assume that the target is located at 0 without loss of generality. Visual feedback of the hand *h*_*t*_ provided by the cursor *c*_*t*_ is given by:

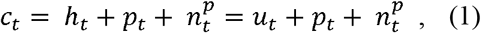

where *p*_*t*_ is the perturbation at time *t*, and 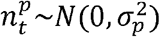 some possible perturbation noise. In order to reach the target at location 0, we assume that a subject generates a motor command *u*_*t*_ to compensate the estimated perturbation 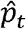, on which a strategy and/or a correction *S*_*t*_ term can be added:

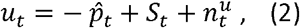

with d 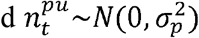 some possible motor noise.

Receiving the efferent copy of the motor command, the internal forward model independently predicts the hand position from its own perturbation estimate:

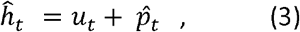

where 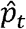 is the perturbation estimate. The SPE is given by:

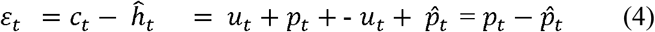

Such SPE can then update the estimate of the perturbation:

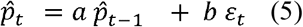

where *a* is a forgetting rate and *b* a learning rate. As can be seen, according to this simple model, the SPE, and memory update, are independent of the actual movements.

Such independence of internal model update from actual motor commands was partially supported in a previous study that studied differences in adaptation in shooting vs. reaching movements ^23^. Whereas in the shooting condition, no motor correction was possible, participants could correct for on-line errors in the reaching condition. No differences in the rate of adaptation were found in healthy control participants, suggesting that neither actual motor movements nor motor errors were responsible for internal model update. However, the effect of the performance errors was unclear in this paradigm since the participants were always exposed to visual targets, which afforded them to compute performance errors. Here, because we manipulated the potential contribution of the targets by eliminating them after movement onset, we showed that SPEs, not performance errors, were the major error signals that updated the forward models.

Despite important differences in experimental procedures, our results of identical washouts in the four conditions, despite large differences in behavior during adaptation, are in line with the results of two other types of arm visuomotor adaptation experiment. First, Miyamoto et al. ^24^ performed a visuomotor adaptation experiment in which participants adapted to a visuomotor perturbation to a single target. There was a remarkable convergence of the stable implicit component of adaptation after a one-minute break across participants. Because there was no relation between the stable component of adaptation and strategy (their figure Fig 3l), it was concluded that the implicit stable component of adaptation is driven by SPEs. Note however the difference between the Miyamoto ^24^ and the current study. Whereas in their study the overall performance during adaptation was highly stereotyped between participants, it was highly variable in the present study. Second, in a type of experiment called “visual clamp”, participants were instructed to perform straight ahead movements while the cursor was rotated to different angles ^25^. Remarkably, the amount of adaptation, as measured by a drift in hand directions was highly similar across a number of cursor rotations. Note here however that it is unclear how SPEs are involved in the drift in visual clamp experiments, since the drift is constant despite largely variable SPEs across cursor rotation conditions.

How can we reconcile the view that reward-related signals play a role in motor adaptation ^8,11–14^? In the present study, no reward was given, thus performance error PE1 could have played the role of an implicit (e.g., self-evaluation) reward. Unlike actual extrinsic rewards, our results show that performance errors do not seem sufficient to influence the update of temporally stable component of motor memories in adaptation, at least in the time scale of the delayed washout period in our static arm reaching experiment. Note, however, that the effect of rewards on adaptation may be more pronounced in dynamic environment than in our static environment ^26^. Another possibility is that performance errors act mostly as punishments in our study, since we do not provide explicit rewards when the cursor is within the target (such as commonly given “target explosions”). Unlike rewards which increase retention ^13,14^, punishments appear to only modulate short-term increase in performance ^13^ and not retention. This is in line with our data because PE1 modulates performance during adaptation, but not performance during washout.

In summary, our results suggest that the temporally stable component of motor memory is formed by SPEs alone, whereas the strategy and the fast process ^24^ can be altered by performance errors during the period of motor adaptation. This differential update of the components of adaptation can account for the large “learning-performance distinction” ^27^, according to which performance during training is a poor predictor of long-term performance. The results of the current study shed further light on the mechanism underlying the learning-performance distinction during learning of motor behaviors and can help with the development of algorithms designed to enhance motor learning in rehabilitation and sports ^28^.

## Acknowledgements

Research reported in this publication was supported by the National Institutes of Health under Award Numbers R01 HD065438 and R56 NS100528. The content is solely the responsibility of the authors and does not necessarily represent the official views of the National Institutes of Health. We thank Cheolhwan Sim for illustrations on Figure 1B, and Annette Eom for comments on our earlier version.

## Author contributions statement

K. L., N. S., and J. I. conceived and designed the experiments. K. L. performed the experiments and analyzed the data. Y. O. provided helps in the experiments. K. L., J. I., and N. S. co-wrote the manuscript. All authors discussed the results and commented on the manuscript.

